# Reconstructing neural representations of tactile space

**DOI:** 10.1101/679241

**Authors:** Luigi Tamè, Raffaele Tucciarelli, Renata Sadibolova, Martin I. Sereno, Matthew R. Longo

## Abstract

Psychophysical experiments have demonstrated large and highly systematic perceptual distortions of tactile space. We investigated the neural basis of tactile space by analyzing activity patterns induced by tactile stimulation of nine points on a 3 × 3 square grid on the hand dorsum using functional magnetic resonance (fMRI). We used a searchlight approach within pre-defined regions of interests (ROIs) to compute the pairwise Euclidean distances between the activity patterns elicited by tactile stimulation. Then, we used multidimensional scaling (MDS) to reconstruct tactile space at the neural level and compare it with skin space at the perceptual level. Our reconstructions of the shape of skin space in contralateral primary somatosensory (SI) and motor (M1) cortices reveal that it is distorted in a way that matches the perceptual shape of skin space. This suggests that early sensorimotor areas are critical to processing tactile space perception.

**Significant Statement:** Here, we show that the primary somatosensory (SI) and motor (M1) cortices, rather than higher-level brain areas, are critical to estimating distances between tactile stimuli on the hand dorsum. By combining functional magnetic resonance (fMRI), Procrustes alignment, and multidimensional scaling, we reconstructed the shape of skin space in the brain. Strikingly, the shape of the skin that we reconstructed from neural data matches the distortions we found at the behavioral level, providing strong evidence that early sensorimotor areas are critical for the construction of tactile space. Our work therefore supports the view that tactile distance perception is computed at lower level in the somatosensory system than is usually supposed.

## Introduction

Perceiving the physical properties of objects through touch is critical for everyday behavior. Since the pioneering work of Weber (1834/1996), the perception of tactile distance has been widely used to investigate the somatosensory system and its links to higher-level aspects of body representation. Recent results have shown that tactile distance is susceptible to sensory adaptation (Calzolari, Azañón, Danvers, Vallar, & Longo, 2017), suggesting that it might be a basic feature coded at relatively early stages of somatosensory processing. Indeed, there is evidence that perceived tactile distance is shaped by low-level features of somatosensory organization such as cortical magnification (Cholewiak, 1999; Taylor-Clarke, Jacobsen, & Haggard, 2004; Weber, 1834/1996) and receptive field (RF) geometry (Brown, Fuchs, & Tapper, 1975; DiCarlo, Johnson, & Hsiao, 1998). Other results, however, show that tactile distance is also modulated by higher-level factors, including tool use (Canzoneri et al., 2013; Miller, Longo, & Saygin, 2014), categorical segmentation of the body into discrete parts (de Vignemont, Majid, Jola, & Haggard, 2009; Le Cornu Knight, Longo, & Bremner, 2014), and illusions of body part size (de Vignemont, Ehrsson, & Haggard, 2005; Tajadura Jimenez et al., 2012; Taylor-Clarke et al., 2004). Together, these results suggest that tactile distance perception is shaped by a combination of bottom-up and top-down factors. The neural bases of this ability, however, remain uncertain.

One source of information about the mechanisms underlying tactile distance perception comes from studies of tactile distance illusions indicating that the representation of the skin surface is systematically distorted. For example, in his seminal work Weber (1834/1996) found that when moving the two points of a compass across the skin, the perceived distance changed, feeling larger on more sensitive skin regions (e.g., the hand) than on less sensitive regions (e.g., the forearm). This effect is known as *Weber’s Illusion*, and subsequent studies have found a systematic relation across the skin between cortical magnification factors and perceptive tactile distance (Cholewiak, 1999; Sadibolova, Tamè, Walsh, & Longo, 2018). This suggests that distortions of primary somatotopic maps, for example of the famous Penfield homunculus (Penfield & Boldrey, 1937), are preserved in some aspects of higher-level tactile perception, therefore, a complete process of tactile size constancy may not always be achieved. Analogous distortions are also found when the same skin region is stimulated in different orientations. For instance, Longo and Haggard (2011) found a bias to overestimate the distance between touches oriented with the medio-lateral hand axis compared to the proximo-distal axis. Similar anisotropies have been reported on several other body parts, including the forearm (Green, 1982; Le Cornu Knight et al., 2014; Marks et al., 1982), thigh (Green, 1982), shin (Stone, Keizer, & Dijkerman, 2018), and face (Fiori & Longo, 2018; Longo, Ghosh, & Yahya, 2015). Intriguingly, such illusions mirror anisotropies in the geometry of tactile RFs which in animals tend to be oval-shaped both in the spinal cord (e.g., Brown, Fuchs, & Tapper, 1975) and in SI (e.g., Alloway, Rosenthal, & Burton, 1989; Brooks, Rudomin, & Slayman, 1961) with the longer axis aligned with the proximo-distal body axis.

The neural mechanisms underlying the perception of tactile distance remain unclear. On one model, perceived distance may be a relatively direct readout of the structure of tactile space as coded by body maps in early somatosensory cortex. This interpretation is supported by the fact that tactile distance adaptation aftereffects show low-level characteristics such as orientation- and location-specificity (Calzolari et al., 2017), as well as by the relation between tactile distance illusions and factors such as cortical magnification and RF geometry. In this case, the representation of the body (i.e., hand) in SI should mirror the distortions observed perceptually. On another model, tactile distance may be calculated at higher-level processing stages such as for instance in the posterior parietal cortex at which distorted primary representations of the skin may be (at least partially) corrected, a form of tactile size constancy. For example, Huang and Sereno (2007) found that the overrepresentation of the lips relative to the rest of the face seen in SI maps is reduced in face maps in the ventral intraparietal area (VIP). This interpretation is supported by: (1) the fact that factors such as illusions of body size and tool use alter perceived tactile distance, which suggests that tactile distance perception is not a direct readout of low-level tactile processing, (2) the fact that while tactile distance illusions mirror distortions of somatotopic maps they are much smaller in magnitude (Longo, 2017; Taylor-Clarke et al., 2004), and (3) the finding that disruption of processing in posterior parietal cortex with transcranial direct current stimulation (tDCS) impairs perception of tactile distance (Spitoni et al., 2013). On this model, the representation of the body in posterior parietal cortex should mirror perception, whereas SI should show much larger distortions – i.e., greater anisotropy.

In this study we investigated the neural mechanisms underlying tactile distance perception by directly comparing neural and perceptual maps of the hand dorsum. We applied a method we recently developed to reconstruct perceptual configurations from the pattern of distance judgments using multidimensional scaling (MDS) (Longo & Golubova, 2017). MDS is a method for reconstructing the latent spatial structure underlying a set of items given a matrix of pairwise distances or dissimilarities between items (Cox & Cox, 2001; Shepard, 1980; for a similar approach applied to neurophysiological data see Sereno & Lehky, 2011). Longo and Golubova obtained judgments of the distance between touches applied to every pair of 16 locations arranged in a 4×4 grid on the hand dorsum. By applying MDS to the resulting perceptual distance matrix, they constructed perceptual maps of the skin which they then compared to actual skin shape. These configurations were clearly distorted, being elongated in the medio-lateral hand axis.

Here, we constructed neural maps of tactile space in an analogous manner. We used representational similarity analysis (Kriegeskorte, Mur, & Bandettini, 2008; Kriegeskorte, Mur, Ruff, et al., 2008) to investigate the structure of tactile space in different brain areas. By applying MDS to the representational dissimilarity matrix for a set of skin locations in a region of interest (ROI), we could reconstruct the neural representation of tactile space, and compare these configurations to the perceptual ones and to actual skin shape.

## Results

### Behavioral data

In an initial behavioral session (N=12 participants), we measured perceptual representation of the skin surface of the hand dorsum using the method developed by Longo and Golubova (2017). An MR-compatible air-puff system (Dodecapus; Huang & Sereno, 2007; see Figure 1A) was used to apply tactile stimulation to nine locations arranged in a 3×3 grid on the dorsum of the participant’s right hand (see Figure 1B). On each trial, two locations were stimulated in sequence and the participant indicated the perceived distance between them by adjusting the length of a visually-presented line on a monitor. By obtaining estimates of every combination of locations, we obtained a perceptual distance matrix of the nine locations for each participant. We then used MDS to construct 2-dimensional perceptual configuration of the skin, as in our recent study using this paradigm (Longo & Golubova, 2017).

**Figure 1.**
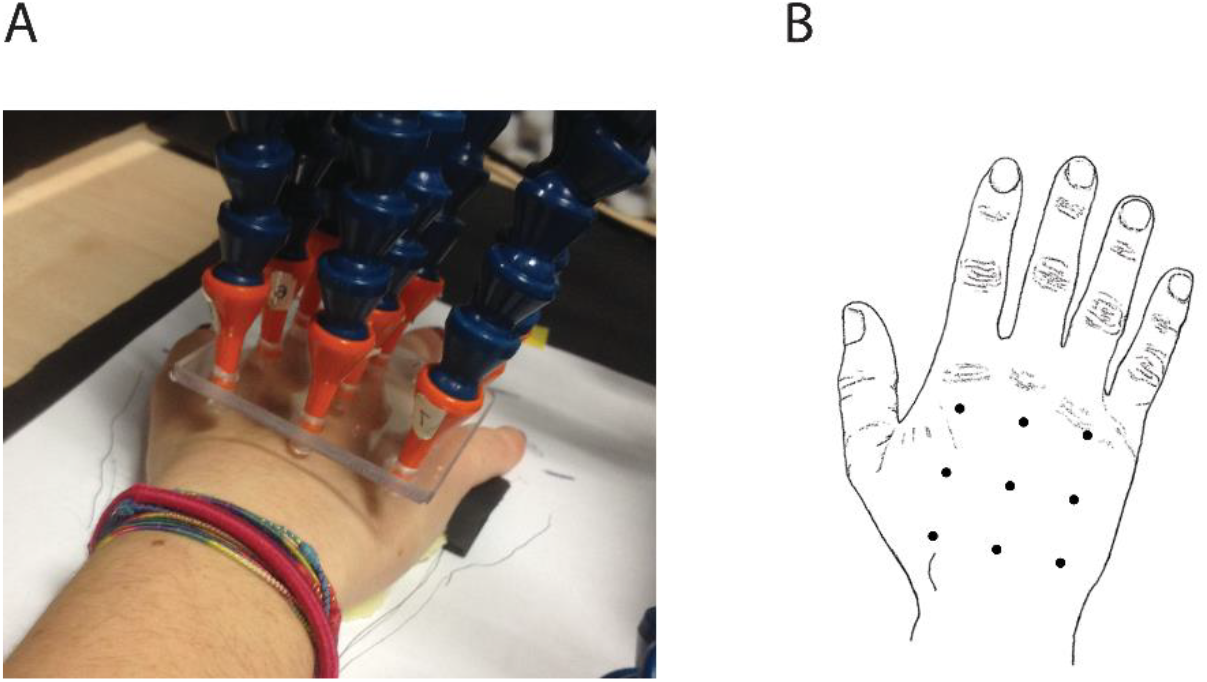
Picture from the participant’s perspective of the apparatus used to deliver air puff stimulation (A). The nine air puff nozzles were positioned on the top of the participant’s right hand dorsum and partially inserted into a plastic plate specifically designed to keep them in place forming a perfect square grid (5×5cm). The six air puff nozzles of the left and right sides of the plate were not perpendicular to the hand, but slightly tilted in the anti-clockwise (left) and clockwise (left) directions in order to resemble the curvature of the hand dorsum. Therefore, all the nozzles were positioned perpendicular to the skin surface. The nozzles were positioned at approximately 3 mm from the skin surface to prevent direct contact with it. Schematic representation of the position of the points on the dorsum of the right hand from a top view (B). Note that vision of the hand was always prevented.

These configurations are shown in Figure 2A (Individual data are shown in Figure S1 of the supplementary material). In order to quantify distortion of these configurations, we estimated the stretch applied to an idealized square 3×3 grid that minimized the dissimilarity with each configuration, as in previous studies (e.g., Longo & Golubova, 2017; Longo & Morcom, 2016). Stretches were defined by multiplying the *x*-coordinates of a square grid (reflecting location in the medio-lateral hand axis) by a *stretch* parameter. Thus, stretch of 1 indicates a perfectly square grid, stretch of less than 1 indicates a tall thin grid, and stretch of greater than 1 indicates a squat fat grid. For each configuration, we identified the value of the stretch parameter that minimized the dissimilarity in shape between the stretched grid and the configuration, quantified as the Procrustes distance between the two configurations. The Procrustes quantifies the dissimiliarity between two configurations as the root mean-square of the residuals after removing differences in translation, rotation, and scale (Bookstein, 1991).

**Figure 2.**
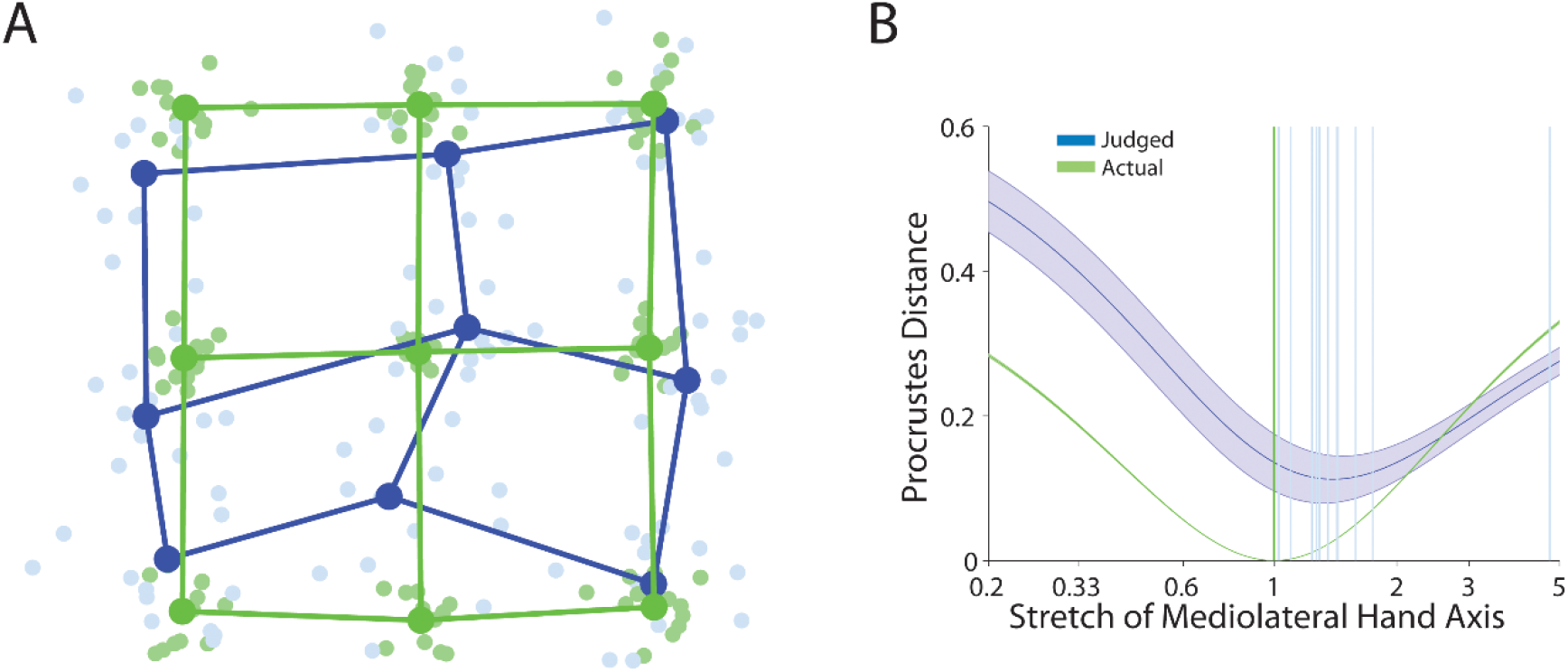
Perceptual hand representation of the spatial configuration of the skin surface. (A) Generalized Procrustes alignment of the actual configuration of points on the hand (green dots and lines) and perceptual configurations (blue dots and lines). The light dots are data from individual participants, while the dark dots represent the averaged shape. (B) Mean Procrustes distance of the perceptual configurations for each participant and idealized grid stretched by different amounts. A stretch of 1 indicates a square grid; stretches greater than 1 indicate stretch in the medio-lateral axis, while stretches less than 1 indicate stretch in the proximo-distal axis. Blue lines represent values for each participant.

Figure 2B shows the mean Procrustes distance for values of the stretch parameter between 0.2 and 5. The best-fitting stretch parameters were significantly greater than 1 (*M* = 1.47), *t*(11) = 3.38, *p* < 0.006, Cohen’s *d* = 0.97, indicating a substantial bias to overestimate distances in the medio-lateral compared to the proximo-distal hand axis. (Note that for this and other tests involving ratios, the calculation of means and all statistical tests were conducted on log-transformed values, which were converted back to ratios to report mean values). As predicted, this result replicates the anisotropy in tactile distance perception previously reported on the hand dorsum (Longo & Golubova, 2017; Longo & Haggard, 2011).

### fMRI data

The main aim of this study was to identify the neural bases of these distorted perceptual spatial configuration of the skin surface. Accordingly, in a subsequent session we analyzed activity patterns using fMRI during tactile stimulation of the same nine points on the hand dorsum. For each participant (the same ones that participated to the behavioral session), we measured the response patterns elicited by each stimulated point estimating the gain parameters (betas) using a general linear model (GLM) approach for each stimulation condition which were used as regressors of interest (see the Materials and Methods section). We then applied representational similarity analysis (RSA; Kriegeskorte, Mur, & Bandettini, 2008) to estimate the geometry of the neural tactile space. More specifically, for a given region of interest, we calculated the Euclidean distance between pairs of neural patterns elicited by the nine stimulus locations. This yielded a representational distance matrix, which we then used to construct 2-dimensional neural configurations of the stimuli using MDS, exactly as we did with the behavioral data. In adopting this approach, we investigated the neural configurations of the hand by investigating similarities between distributed patterns of activity, rather than by measuring non-overlapping, somatopically organized foci in a somatotopic representation (cf. Ejaz, Hamada, & Diedrichsen, 2015).

We investigated neural configurations of the hand in several ROIs known to be involved in the processing of tactile stimuli, including primary (SI) and secondary (SII) somatosensory cortex, primary motor cortex (M1), and the superior parietal lobe (SPL), all in the contralateral left hemisphere. As control ROIs, we used the early visual cortex (EVC), as well as all the same ROIs in right hemisphere ipsilateral to the locus of stimulation (for ROIs specification see the Materials and Methods section). Within each ROI, we used a searchlight procedure to identify brain areas from which shape of the skin could be successfully reconstructed. For each searchlight, we calculated the Procrustes distance between the resulting neural configuration and either the participant’s perceptual configuration or the actual structure of the skin.

Figure 3 shows the topographic distribution of the resulting group Procrustes distances between the neural and perceptual configurations. Warm colors indicate small Procrustes distance, and thus high similarity between the shapes. To identify clusters of Procrustes distances statistically smaller than the chance level, we adopted a bootstrapping procedure as suggested by Stelzer, Chen, and Turner (2013). In brief, for each participant and ROI, we computed again the Procrustes distances as explained above except that we shuffled the labels of the stimulated conditions before computing the Euclidean distance. We thus obtained 100 random Procrustes distance maps and we then used the bootstrapping procedure (10,000 iterations) to estimate the distribution of cluster sizes under the null hypothesis. Clusters in the actual data with a size exceeding the critical size value associated with a p-value of 0.05 were considered significant (see the Materials and Methods section for a more detailed description of the procedure). Procrustes distances significantly smaller than chance (see Figure 3B for the histograms of the Bootstrap values for the brain regions of the pre-defined ROIs in which we were able to reconstruct the geometry of the skin) and Figures S5 and S6 in the supplementary materials for the other brain regions) were found in clusters observed in contralateral SI (one in area 3b/1 and another in area 2) and M1 (area 4) only, as shown in Figure 3, Panel A and C (red contours indicate the significant clusters). Table 1 reports the outcome of the cluster analysis. No significant clusters were observed in the EVC or in the ipsilateral ROIs. Nearly identical results were obtained when we compared neural configurations to the actual grid shape (see Figure S5 of the supplementary materials). These results show that the perceptual structure of the skin can be reconstructed from the representational pattern in both primary somatosensory and motor cortices in the contralateral hemisphere. The shapes associated with each significant cluster are shown in Figure 3D superimposed on the behavioral and actual shapes.

**Table 1.**
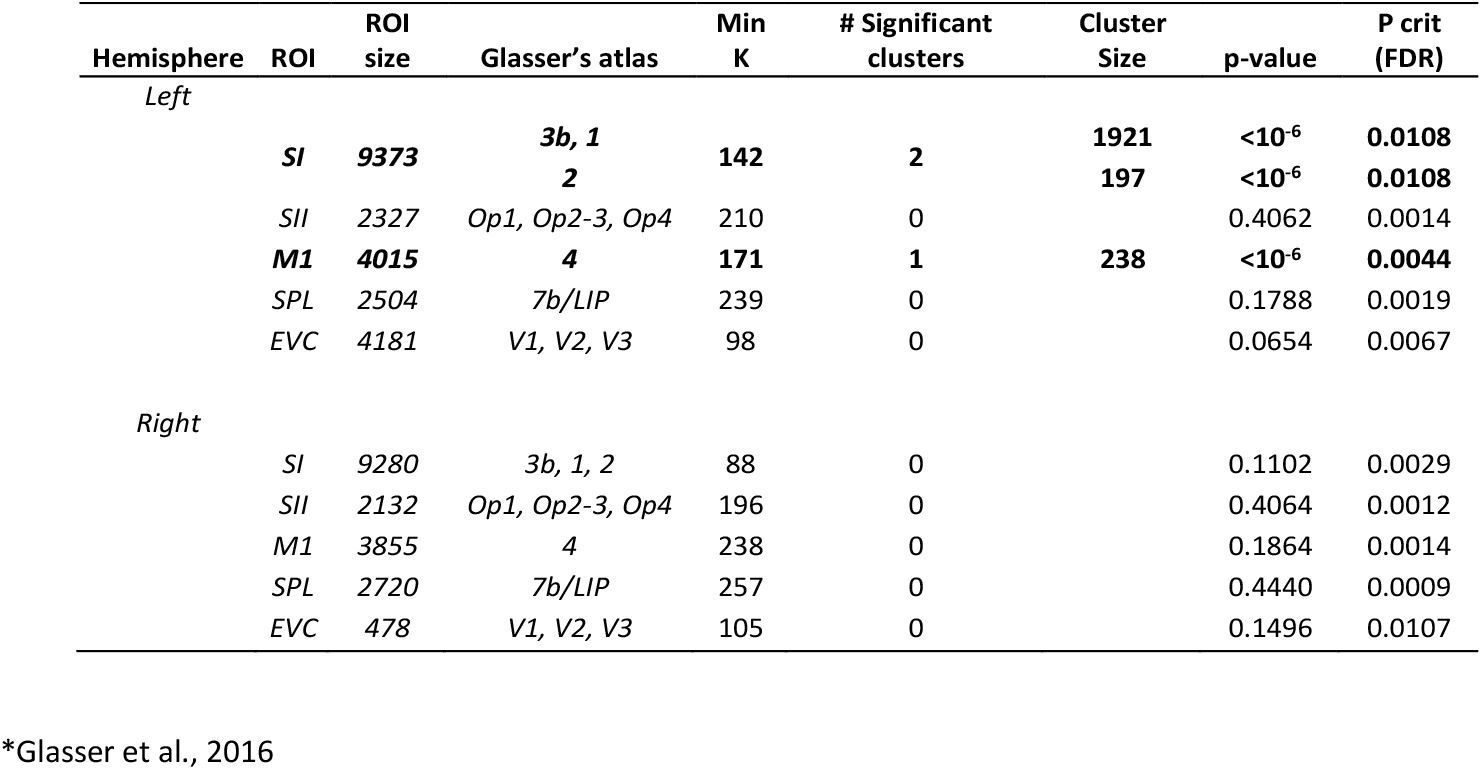
For each ROI, we show the minimum dimension (**Min K**) a cluster should have to be considered significant as resulting from the cluster-based bootstrapping analysis (p<0.001 at the vertex level; FDR<0.05 at the cluster level). For the significant clusters only (in bold), the size is also reported (rightmost column).

**Figure 3:**
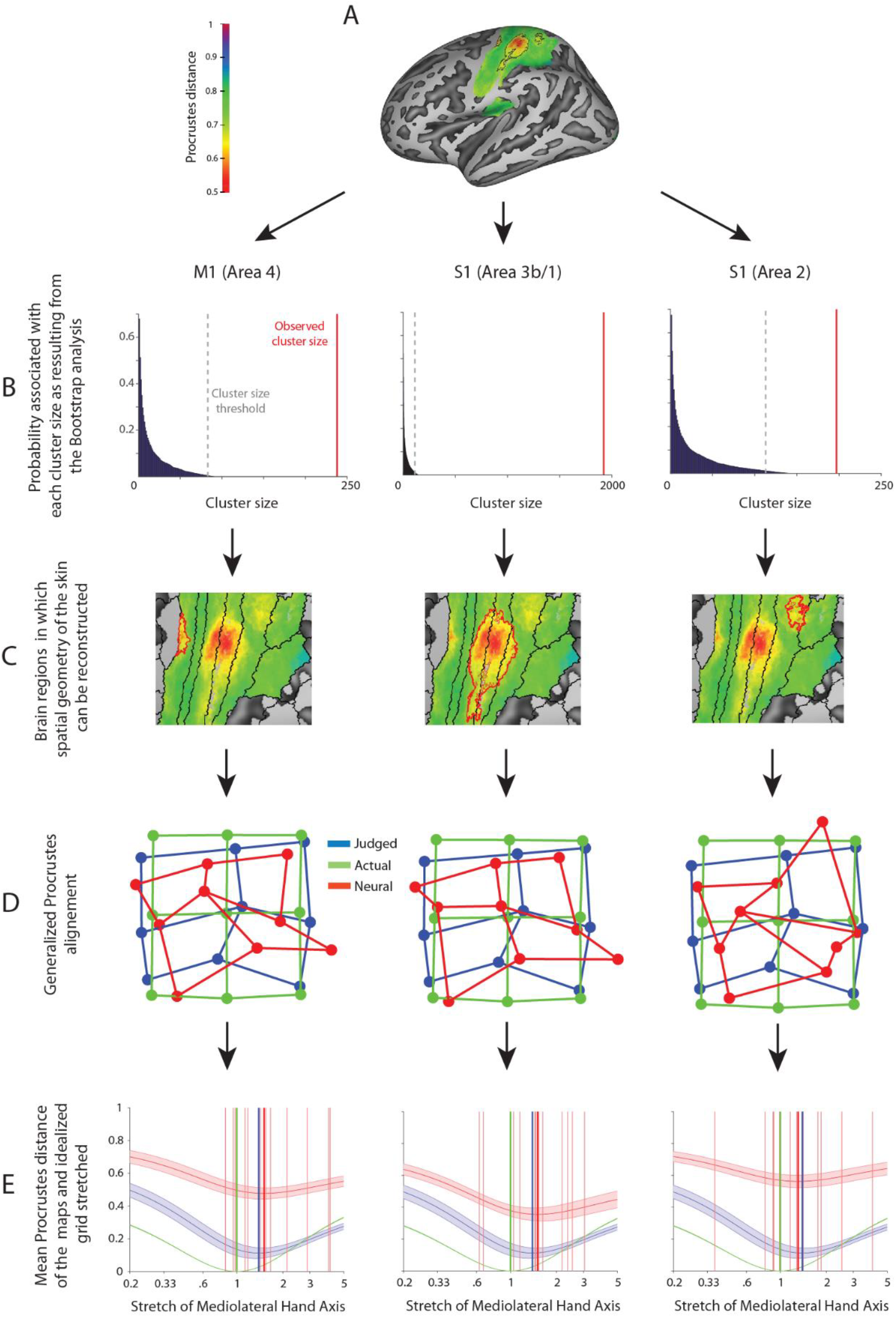
(A) Brain regions in which the spatial geometry of the skin could be reconstructed from the representational pattern of neural activations. Red contours reflect significant cluster resulted from the cluster-based bootstrapping analysis (p<0.001 at the vertex level; FDR<0.05 at the cluster level). (B) Probability associated with each cluster size as resulting from the Bootstrap analysis. The dotted gray lines represent the critical size values (cluster p-value<=0.05) for each ROI; the red lines represent the actual size of the observed cluster. Only the significant clusters are shown in this figure, refer to the supplementary Material for the other ROIs (C) Magnified view of the three significant clusters for area 4 (M1), area 3b/1 (SI) and area 2 (SI), respectively, when comparing the neural and perceptual configurations. The red color represents the voxels in which the reconstructed representations were better achieved as expressed in Procrustes distance value. (D) Generalized Procrustes alignment of the grand average shape across participants of the actual configuration of points on the dorsum of the right hand (green dots and lines), perceptual (blue dots and lines) and neural (red dots and lines) configurations. (E) Mean Procrustes distance between behavioral (blue), fMRI (red: for each participant) and actual (green) grid on the participants’ hand dorsum and idealized grids stretched by different amounts. A stretch of 1 indicates a square grid; stretches greater than 1 indicate stretch in the medio-lateral axis, while stretches less than 1 indicate stretch in the proximo-distal axis. The shaded regions indicate one standard error of the mean. The dotted vertical lines indicate the mean of the best-fitting stretches for fMRI (red) and behavioral configurations (blu). The stretch that minimized the Procrustes distance was substantially larger than 1. Thus, there was clear evidence for stretch in the medio-lateral hand axis for perceptual configurations.

We investigated distortions at the level of neural representations using the same analysis of stretch applied to the behavioral data above. Figure 3E shows the Procrustes distances as a function of the stretch for the perceptual, actual, and fMRI configurations (Procrustes distances as a function of the stretch for each individual participant are reported in Figures S2, S3 and S4 in the supplementary material).

Moreover, we also computed the Procrustes distance analysis at the whole brain level by performing a cluster-based bootstrapping analysis (p<0.001 at the vertex level; FDR<0.05 at the cluster level) on the whole brain to identify potentially significant clusters beyond the pre-defined ROIs. Such analysis confirmed the resulted significant clusters performed on the pre-specified ROIs, moreover, some other cluster resulted to be significant (see Figure S7 in the supplementary material), namely in the contralateral hemisphere Area 55b and OP4 (SII) and in the ipsilateral hemisphere what we define as the parietal operculum (PO) (note that the PO cluster was positioned in the straddling areas 2, PFt and PFop) and the Superior Temporal Visual area (STV).

Despite the presence of additional significant clusters the quality of the reconstruction of the spatial geometry was not satisfactory for such clusters (Figure S8 in the supplementary material). However, despite its lowers Procrustes distance value the reconstructed configuration for area PO was good resembling the shape of the original undistorted grid.

Best-fitting values of the stretch parameter were significantly greater than 1 for the cluster in SI straddling areas 3b and 1 (*M*: 1.47), *t*(11) = 2.68, *p* = 0.021, *d* = 0.77, and the cluster in M1 (*M:* 1.64), *t*(11) = 3.14, *p* = 0.010, *d* = 0.91. However, for the cluster within area 2 of SI this distortion was not significant (*M*: 1.26), *t*(11) = 1.33, *p* = 0.212, *d* = 0.38. Moreover, none of these clusters showed a significant stretch parameter different from those observed for the perceptual configurations given by the behavioral data (all *p* > 0.45). Virtually identical results were obtained for the clusters identified by comparing neural configurations to actual skin configuration shape (see Figure S5 in the supplementary materials).

## Discussion

In the present study, we reconstructed the internal geometry of tactile space using representational similarity of neural patterns between locations on the skin. Behaviorally, we replicated previous reports that tactile space is stretched along the medio-lateral axis of the hand dorsum (e.g., Longo & Golubova, 2017; Longo & Haggard, 2010, 2012). Critically, using a novel approach that combine fMRI with MDS, we showed that similar distortions can be measured directly from neural data. Strikingly, this was evident in the primary sensorimotor cortices contralateral to the locus of stimulation. Therefore, these low-level cortical brain areas carry information corresponding to the distorted perceptual structure of tactile space of the hand dorsum being stretched along the medio-lateral axis.

Interestingly, the sensorimotor cortices were the only brain areas from which we were able to reconstruct maps of the shape of the skin. Previous studies have shown the presence of clear somatotopically organized representations of different body parts in SI contralateral to the locus of stimulation (e.g., Huang, Chen, Tran, Holstein, & Sereno, 2012; Sanchez-Panchuelo, Francis, Bowtell, & Schluppeck, 2010). However, this does not seem to be the case for the hand dorsum, in which, to the best of our knowledge, clear maps have not been shown in humans. Recently, a study has shown only a difference in terms of peak of cortical activation and numbers of activated voxels between the dorsum and the palm of the hand, with the former being lower than the latter (Jang, Seo, Ahn, & Lee, 2013). Moreover, in the monkey neurophysiological literature, it is unclear to what extent similar topographic maps (i.e., palm and dorsum of the hand) can be clearly defined, given that these neurophysiological studies have shown that representations of the dorsal hand surface may fall outside the global somatotopic pattern in SI (Kaas, 1983).

The distortions of the neural maps we constructed from representational similarity of neural patterns in contralateral sensorimotor cortex provide an intriguing correspondence with the anisotropic geometry of (RFs) in the somatosensory cortex (Alloway et al., 1989; Brooks et al., 1961). We have proposed that tactile space can be thought of as a 2-dimensional array in which the RFs of neurons in somatotopic maps forming the “pixels” of the grid (Fiori & Longo, 2018; Longo, 2017; Longo & Haggard, 2011). Where RFs differ in size on different skin surfaces, this will produce a perceptual magnification on the surface with smaller RFs, assuming that RF overlap is comparable (that is, assuming that regions with smaller RFs occupy proportionally more cortical area). Neurophysiological studies have provided some evidence for this assumption, finding that overlap between the RFs of adjacent neurons is a constant proportion of RF size across a wide range of sizes (Sur, Merzenich, & Kaas, 1980). Where individual RFs are anisotropic (e.g., oval-shaped), this will produce a perceptual stretch along the shorter axis of the RF. The somatosensory RFs on the hairy skin of the limbs tend to be oval-shaped with the long-axis aligned with the proximo-distal limb axis (e.g., Brooks et al. 1961; Alloway et al. 1989), compatible with the results of the present study. However, the magnitude of these distortions is much smaller than what would be predicted only on the basis of differences in RF size and shape. Indeed, the long axis of RFs in somatosensory cortex is frequently 4 –5 times the length of the small axis (e.g., Brooks et al., 1961), yet the magnitude of perceptual anisotropy is again only a small fraction of that (e.g., Green, 1982; Longo & Golubova, 2017; Longo & Haggard, 2011b). We suggest that a process of tactile size constancy which corrects for distortions inherent in primary representations to produce (approximately) veridical percepts of size may take place in the sensorimotor cortices, particularly, in the primary somatosensory cortex where the reconstructed skin shape was more accurate. In agreement with our results, a recent study using repetitive Transcranial Magnetic Stimulation (rTMS), have shown that the metric representation of the body depends on somatosensory afferences. In their study Giurgola and colleagues (2019) applied rTMS on the somatosensory cortices of both hemispheres representing the hands (i.e., SI) while participants judge whether visually presented right and left hands matched the size of their own hand. They found that rTMS produces distortions of the perceived size of the participants’ own hand, but not other body parts (Giurgola, Pisoni, Maravita, Vallar, & Bolognini, 2019). Intriguingly, this effect was not present when rTMS was applied on the inferior temporal parietal lobe, an area largely linked with body representation disturbances (Bolognini & Miniussi, 2018). However, our approach did not allow us to rule out possible top-down interactions between SI and other brain areas (e.g., higher level regions), thus preventing any definite conclusion about the pathway leading to our results. Indeed, other brain areas may have interacted with the primary somatosensory cortex (and/or primary motor cortex) providing information to correct for homuncular distortions. It would be relevant to assess this question in a dedicated study which possibly involve other neuroimaging techniques with a higher temporal resolution than fMRI such as electroencephalography (EEG) or magnetoencephalography (MEG).

These results support the notion that the computation of distance perception between tactile points on the skin of the hand dorsum is computed at a low cortical level of tactile representation processing. In this respect, Calzolari and colleagues (2017) using a tactile adaptation aftereffect paradigm suggested that tactile distance perception is a basic somatosensory feature supporting the idea that distance perception arises at relatively early stages in tactile processing. In their study, the authors explored how adaptation to a distance between two separate points, passively delivered on the hand dorsum, affects perception of subsequent distances. They found tactile distance aftereffects with passive touch. Moreover, their effect was orientation and region specific, did not transfer within and between the hands, and was encoded using skin-based coordinates. These are all features that point to a low level processing locus for tactile distance computation. Similarly in vision, Sperandio and colleagues (2012) found that the retinotopic activity in the primary visual cortex (V1) reflects the perceived rather than the retinal size of an afterimage. The fact that SI is critically involved in a complex processing such as tactile distance estimation is in accordance with literature showing that this low cortical level area may not be critical for performing simple tactile tasks – i.e., tactile detection – both in monkeys (LaMotte & Mountcastle, 1979) and humans (Tamè & Holmes, 2016). By contrast, SI seems to be critically involved in processing that were thought to be accomplished by higher level cortical areas, such as bilateral integration of touch (Tamè, Braun, Holmes, Farnè, & Pavani, 2016; Tamè et al., 2012; Tamè, Pavani, Papadelis, Farnè, & Braun, 2015) as well as tactile working memory (Harris, Miniussi, Harris, & Diamond, 2002; Katus, Grubert, & Eimer, 2015).

Other behavioral studies that used a different paradigm which investigated participants’ abilities to localize the position of the different parts of the hand relative to each other showed the presence of similar distortions. In this respect, Longo and Haggard (Longo & Haggard, 2010, 2012) asked participants to place their hand flat on a table underneath an occluding board and to use a long baton to judge the perceived location of the tip and knuckle of each of their finger. By comparing the relative location of judgments of each landmark, authors constructed perceptual configurations of hand structure which they then compared to actual hand form. A highly consistent pattern of distortions was apparent across participants, including overestimation of hand width, and underestimation of finger length. Longo, Mancini, and Haggard (2015) conducted a similar study, but asked participants to judge the location of tactile stimuli applied to the hand dorsum, finding overestimation of distances in the medio-lateral hand axis, compared to the proximo-distal axis. Interestingly, this pattern of distortions is quite similar to that described in the present study. Therefore, the present results further support the idea that similar mechanisms may underlie body position sense and tactile distance perception (Longo & Haggard, 2010).

The fact that we were able to reconstruct the shape of the skin space based on activation elicited by tactile points both in the primary somatosensory and motor cortices suggests that M1 is also involved in the processing of the tactile stimuli. In everyday life, tactile stimulation is commonly accompanied or caused by action. Indeed, the sensory and motor systems are intimately related, both anatomically and functionally, with continuous reciprocal exchange of information (Brochier, Boudreau, Paré, & Smith, 1999; Nelson, Staines, & McIlroy, 2004; Rossi, Pasqualetti, Tecchio, Sabato, & Rossini, 1998). These systems communicate via a network of extensive connections between the sensory and motor cortices (Andersson, 1995; Asanuma, Stoney, & Abzug, 1968; Eickhoff et al., 2010; Huffman, 2001; Makris et al., 2005; Mao et al., 2011; Shinoura et al., 2005; Stepniewska, Preuss, & Kaas, 1993; Strick & Preston, 1982), but also by motor cortex cells responding directly to sensory stimuli, perhaps via their direct inputs from the dorsal column nuclei via the ventrolateral thalamic nucleus (Albe-Fessard & Liebeskind, 1966; Fetz, Finocchio, Baker, & Soso, 1980; Fromm, Wise, & Evarts, 1984; Goldring & Ratcheson, 1972) and vice-versa (Matyas et al., 2010). The existence of direct connections between the sensory areas in the post-central gyrus and the motor areas of the precentral gyrus in humans has been recently demonstrated by Catani and colleagues who, using diffusion tractography, confirmed the presence of U-shape fibres that directly connect SI with the motor cortex (Catani et al., 2012), as previously demonstrated in invasive studies in animals. These fibers are thought to connect the somatosensory and motor areas of the cortical regions that are involved in the control of finely tuned movements and complex motor skills (i.e. the hand’s brain regions). In this respect, Tamè, Pavani, Braun, Salemme, Farnè and Reilly (2015) combined tactile repetition suppression with the techniques of afferent inhibition (i.e., corticospinal excitability is inhibited when a single tactile stimulus is presented before a TMS pulse over the motor cortex) to investigate whether the modulation of somatosensory activity induced by double tactile stimulation propagates to motor cortex and alters corticospinal excitability in humans. They found that activity in the somatosensory cortices following repetitive (i.e., double) tactile stimulation also elicits finger-specific activation in the primary motor cortex demonstrating that spatial information is retained in the SI and then transferred to the motor cortex (Tamè, Pavani, Braun, et al., 2015). Furthermore, the relation between the sensory and motor systems is particularly important in haptic tasks, in which we actively explore an object. In this situation, our brain is simultaneously receiving sensory signals from, and generating motor signals for, the movements. These inputs have to be combined to perceive and actively explored objects. In this respect, Ejaz and colleagues (2015), analyzing activity patterns during individual fingers movements using fMRI, showed that hand use can shape fingers’ arrangement in both the sensory and motor cortices (Ejaz et al., 2015).

Finally, regarding the reconstructed configuration of the tactile space that emerged from the whole brain analysis in the ipsilateral parietal operculum (Figure S8 in the supplementary material) we do not have a definitive interpretation given that this was an unexpected result. A possibility could be that such area is actually currying information about the actual configuration of the skin space, however, such result should be treated with caution given that despite the satisfactory reconstruction, the Procrustes value which represents the difference between the neural and behavioral shapes, was higher than every other brain area.

### Conclusion

In the present study, by applying an innovative approach that combined MDS and Procrustes alignment on fMRI data, we were able to reconstruct the shape of the internal geometry of the skin of the hand dorsum. We showed that the superficial structure of the skin can be reconstructed from the matrix of perceived tactile stimulated points on the hand. Intriguingly, the reconstructed shape of the skin in the primary somatosensory and motor cortices matches the distortions that emerge at behavioral level (i.e., perceptual configurations) providing evidence that sensory-motor cortices may be a primary neural basis of such representations. Intriguingly, the sensorimotor cortices were the only regions that contained sufficiently coherent information to allow a satisfactory reconstruction of the shape of the skin space; we found nothing similar in data from higher level brain regions. We suggest that representations in SI and M1 are likely to be critical for haptic control (Johansson & Flanagan, 2009) of complex hand–object interactions involving events that are precisely localized in space.

## Materials and Methods

### Participants

The same twelve participants (mean ± SD = 29.5 ± 6.3 years; 6 females) participated both in the behavioral and fMRI experiments. Participants reported normal or corrected to normal vision and normal touch. All participants but one were right-handed as assessed by the Edinburgh Handedness Inventory (Oldfield, 1971; M=66 range-90-100). All procedures were approved by the Department of Psychological Sciences Research Ethics Committee at Birkbeck, University of London. The study was conducted in accordance with the principles of the Declaration of Helsinki.

### Air puff stimulation

The MRI-compatible air-puff stimulator is shown in Figure 1A. It was driven by an air compressor in the scanner control room which provided the input to a 9-way solenoid manifold valve (“S” Series Valve; Numatics Inc., Highland, MI) that was controlled by transistor-transistor logic pulses. Nine plastic air tubes from the manifold valve passed through waveguides into the scanner room, where they connected to a block mounted beside the right hand, at the edge of the bore. The block served as a rigid base for 9 flexible tubes with nozzles (Loc-Line Inc., Lake Oswego, OR), flexibly arranged to direct 50 ms air puffs (input air-pressure 3.5 bar) at 9 locations arranged to form a 3×3 grid approximately centered on the dorsum of the right hand (Figure 1A). The tubes were not in contact with the skin surface of the dorsum of the participant’s right hand. Each air puff was perceived as a well-localized and light touch on a specific hand dorsum location.

### Behavioral experiment

#### Procedure

Before the fMRI experiment, participants completed a behavioral experiment in which we asked them to estimate the distance between two touches on the dorsum of their hand. The rationale for this approach was twofold. First, we wanted to test the suitability and effectiveness of our paradigm using the air puff stimulator, which has not previously been used for tasks involving distance judgments. Second, we wanted to have an estimation of the perceptual configuration of the skin in each participant to compare with the neural data. Participants sat comfortably in front of a computer screen, with their right hand lying flat on the table, palm down, with the wrist straight. A black curtain occluded their right hand and forearm. On each trial, participants looked at a black screen and received two sequential air puff stimulations. Each stimulus was delivered to one of the 9 locations on the hairy skin of the hand shown in Figure 1B. Stimulus locations were formed by a 3×3 grid approximately centred on the dorsum. The locations were marked with a felt pen at the start of the study by placing a plastic stencil over the skin surface. Each stimulus consisted of a train of five 50 ms on periods alternating with 50 ms off periods, for a duration of 450 ms. There was a 50 ms inter-stimulus interval between stimulation of the two locations.

Shortly after the second stimulus (jittered randomly between 1-2 s), a line appeared at the center of the screen. The participant was required to adjust the length of the line to match the perceived distance between the two tactile stimuli. There were four possible starting line conditions that occurred in a randomised order and differed by their orientation (horizontal, vertical) and starting length (small: 40 pixels/1.54 cm; large: 460 pixels/17.69 cm). Lines were approximately 1 mm thick and were white on a black background. Participants made unspeeded responses, adjusting the line length on the screen by pressing two arrow buttons on a keypad with the left hand. When they were satisfied with their response, they pressed a third button to confirm their response. Participants were never allowed to look at either hand during the experiment.

There were four blocks with 72 trials each, for a total of 288 trials. In each block, there were 36 possible combinations the 9 points, crossed with two orders of stimulation, which were presented in a random order. At the end of the experiment, a photograph was taken of the participant’s right hand to calculate the actual size of the grid. A ruler appeared in the photographs allowing conversion between distances in pixels and cm. Participants were allowed short breaks between blocks. The experimenter remained in the room throughout the session to ensure that participants complied with the instructions and to keep the position of the hand in place.

#### Multidimensional scaling

Analysis procedures were similar to those in our previous study using this paradigm (Longo & Golubova, 2017). The eight repetitions of each stimulus pair for an individual participant were averaged, resulting in a symmetric matrix reflecting the pairwise perceived distance between pairs of points, with zeros on the diagonal. Classical multidimensional scaling was applied to the distance matrix for each participant using the *cmdscale* command in MATLAB (Mathworks, Natick, MA). The output of MDS is a set of eigenvalues for each dimension and coordinates for each landmark in each dimension. As there are 9 landmarks, MDS attempts to position the landmarks in 9-dimensional space such that the distances between them are as proportional as possible to the perceived distances. To calculate the percentage of variance in the data accounted for by each dimension, we compared the absolute value of each eigenvalue to the sum of the absolute values of all 9 eigenvalues.

In order to create a null distribution for comparison with our data, we conducted MDS on simulated random data. For each simulation, 36 random numbers were generated and placed into a distance matrix, as with the actual data. MDS was applied to each simulation and the eigenvalues and coordinates extracted. One million such simulations were conducted.

#### Procrustes Alignment

Procrustes alignment (Goodall, 1991; Rohlf & Slice, 1990) superimposes two spatial configurations of homologous landmarks by translating, scaling, and rotating them to be as closely aligned as possible. First, the two configurations are translated so that their centroids (i.e., the centre of mass of all landmarks) are in the same location. Second, the configurations are normalized in size so that the centroid size, which is quantified as the square root of the sum of squared distances between each landmark and the centroid, is equal to 1. Third, the configurations are rotated to minimize the sum of squared distance between pairs of homologous landmarks. Note that in the present study mirror reflections of the configurations were allowed, though in other contexts this may not be desirable. At this point, the configurations are in the best possible spatial alignment, with all non-shape differences removed (Bookstein, 1991). We used Procrustes alignment in two ways, both as a way to quantify dissimilarity in shape and as a visualization tool. First, the residual sum of squared distances between pairs of homologous landmarks which is not removed by Procrustes alignment provides a measure of the dissimilarity in shape between the two configurations, called the Procrustes Distance. If two configurations have exactly the same shape, they will lie on top of each other following Procrustes alignment and thus have a Procrustes distance of 0. In contrast, two configurations with no shared spatial structure at all will have a Procrustes distance of 1, given that the size normalization results in a total sum of squared variance within each configuration of exactly 1. Second, Procrustes alignment provides a natural way to visually display configurations, making differences in shape clearly apparent. Given that we had to compare several hand configurations, we used generalized Procrustes analysis (GPA) using Shape (a MATLAB toolbox from Dr Simon Preston, freely available from download [https://www.maths.nottingham.ac.uk/personal/spp/shape.php] based on an algorithm originating from Gower, 1975; TenBerge, 1977).

Finally, we used the Procrustes distance, the sum-of-squares of the residual distances between pairs of homologous landmarks, as a measure of the dissimilarity between two configurations. This allowed us to estimate the overall stretch of perceptual configurations in the medio-lateral axis by finding the stretch applied to an idealized rectangular grid that minimized the dissimilarity with each configuration. We multiplied the x-coordinates of a 3 × 3 square grid by a stretch parameter to generate grids of varying levels of stretch. When the stretch parameter was equal to 1, the grid was perfectly square. When it was greater than 1, the grid was stretched in the medio-lateral axis. When it was less than 1, the grid was stretched in the proximo-distal axis. Note that because Procrustes alignment normalizes size, a stretch applied to the medio-lateral axis is identical to the inverse stretch being applied to the proximo-distal axis. Thus, while distortions are described in terms of the mediolateral axis, this method cannot indicate which specific axis is affected by distortions in the sense that stretch of one axis is formally identical to compression of the other. For each participant, we determined the value of the stretch parameter that minimized the dissimilarity in shape (i.e., that minimized the Procrustes distance) between the stretched grid and the participant’s perceptual configuration. Values between 0.2 and 5 were tested by exhaustive search with a resolution of 0.0005 units in natural logarithm space (i.e., 6,438 steps). Note that we report mean stretch values as ratios, the statistical tests we report compare the mean logarithm of the ratios to 0, since ratios are not symmetrical around 1.

### fMRI experiment

#### Procedure for the main experiment

Participants laid in the scanner with their right hand prone outside the scanner bore, and wore earplugs throughout the experiment. The air-puff stimulators were positioned just over the dorsum of the participants’ right hand as in the behavioral experiment by means of an fMRI compatible plastic plate to arrange the stimulators into a 3×3 grid suspended over the hand without touching the skin (as for the behavioral experiment; see the Results section and Figure 1). At the beginning of each run, participants were instructed to close their eyes and focus their attention on the dorsum of their right hand. Air-puff stimuli were delivered sequentially in a random order on the different 9 points. Each run lasted about 11 minutes and included 55 trials. In each trial, the same point of the skin was stimulated by delivering the air quickly alternating between ON (50ms) and OFF (50ms), except for the asynchronous stimulation that was ON (20ms) and OFF (80ms). The asynchronous stimulation was delivered to ensure that participants were focusing on their right hand as they were asked to report the number of asynchronous stimulations at the end of the run. There were four asynchronous trials per run, for a total of 16 trials in the whole experiment.

Each point was stimulated 5 times in each run for a duration of 12 s. In addition, 10 12-s trials of no stimulation (null trials) were randomly interleaved with the experimental trials.

#### Stimulation and procedure for the functional localiser

After the main experiment participants underwent a functional hand dorsum localizer. Somatosensory stimulation was applied to the dorsum of the right hand, the same location as for the main experiment. Stimulation was performed manually by the experimenter by brushing the participants’ skin with a toothbrush across different directions - i.e., along the proximo-distal and medio-lateral axis, in a back-and-forth manner with a frequency of about 2 Hz. This method has been previously successfully used and proved to be effective in evoking activity in the somatosensory cortices (Disbrow, Roberts, & Krubitzer, 2000; Eickhoff, Grefkes, Zilles, & Fink, 2007). The paradigm consisted of 8 cycles each of them characterised by 16s of stimulation and 16s of rest. The localizer lasted overall 4 minutes.

#### Data acquisition

Echoplanar images (2.33 × 2.33 mm^2^ in-plane, 2.3-mm-thick slices, 662 volumes per run, 36 axial slices, flip = 90°, TE = 39 ms, TR = 1 s, 64 × 64 matrix, bandwidth = 1474 Hz/pixel, data acquired with prospective motion correction) were collected during 4 runs on a Siemens Avanto 1.5 T MRI scanner with a 32-channel head coil. For the functional localizer, echoplanar image parameters were the same as for the main experiment except for the number of volumes that were 256. For the anatomical image, we used an MPRAGE scan (1 × 1 × 1 mm, flip = 7°, TR = 1 s, TI = 1 s, TE = 3.57 ms, matrix 256 × 224 × 176 190 Hz/pixel).

#### Preprocessing and GLM analysis

Before analysis, the first eight volumes of the functional data of each run were discarded to avoid T1 saturation. The anatomical data were segmented using the standard procedure in FreeSurfer (function *recon-all*; Fischl, Sereno, & Dale, 1999), whereas the functional data were preprocessed and analyzed using Statistical Parametric Mapping software (SPM12; Wellcome Centre for Neuroimaging, University College London, London, UK; http://www.fil.ion.ucl.ac.uk/spm). Each functional volume was first bias corrected and then spatially realigned to the first volume of the first run to correct for head movements. The functional volumes were then coregistered to the volumetric anatomical image which was aligned with the surfaces obtained from Freesurfer. First-level analyses were first carried out in the subject space and then the data were normalized to the Freesurfer common space (*fsaverage*). Data were spatially smoothed using a spatial Gaussian kernel of FWHM of 5mm for the univariate second-level analyses only. The multivariate analyses were conducted using the unsmoothed data by means of the Matlab toolbox CoSMoMVPA (Oosterhof, Connolly, & Haxby, 2016) and home-made Matlab scripts.

For each voxel, we estimated the response to the stimulated points by fitting a general linear model (GLM) to the functional data. Each event was modelled using a square-wave function that was convolved with the canonical hemodynamic response. Therefore, for each run, the design matrix was 662 volumes X 9 predictors of interest. We also added six columns to account for head movements and one constant column. The GLM analysis returned 9 betas per run for each voxel, thus we obtained 36 betas of interests. The betas associated with the various points were averaged across runs and the resulted 9 averaged betas were used for the subsequent representational similarity analysis (RSA).

We also ran another similar GLM analysis using the smoothed data and the estimated betas were then used for the second-level univariate analyses. To this aim, we also estimated the betas and t maps associated with the contrast *all stimulated points vs baseline* that were subsequently used for the second-level analysis.

#### Preprocessing and GLM functional localiser

Data preprocessing steps for the functional localiser were identical to the ones performed for the main experiment. For the GLM a single event was modelled using a square-wave function that was convolved with the canonical hemodynamic response. The design matrix was 256 volumes X 1 predictor of interest. We also added six columns to account for head movements and one constant column. The GLM analysis returned 1 betas for each voxel.

#### Identification of ROIs

We identified four regions of interest (SI, SII, M1, SPL) and one as a control (EVC) on the basis of both anatomical and functional criteria (Cavina-Pratesi et al., 2010; Dinstein, Gardner, Jazayeri, & Heeger, 2008; Gallivan, McLean, Valyear, Pettypiece, & Culham, 2011). To create the masks at the surface level, we used the recent multimodal cortical parcellation of the human brain developed by Glasser and colleagues (2016). First, we superimposed the functional localizer with the main experiment functional maps to determine the brain regions that were involved in tactile processing for both type of stimulations. Subsequently, we selected the vertices of interest for each participant based on Glasser and colleague’s parcellation atlas at individual brain space. Specifically, our SI included areas 3a, 3b, 1 and 2 of Glasser and colleagues’ nomenclature; SII included OP1, OP2-3 and OP4; M1 included area 4; SPL included 7PC and AIP. We also analyzed the early visual cortex including V1, V2 and V3 to assess the response pattern also in brain areas outside the typical tactile network. The mean average number of vertices across participants for each ROI are reported in Table 1.

#### Searchlight analysis

We used a searchlight analysis (Kriegeskorte, Formisano, Sorger, & Goebel, 2007; Kriegeskorte, Goebel, & Bandettini, 2006) to identify the brain regions that contain meaningful activity patterns about the spatial configuration - i.e. shape - of the skin of the dorsum of the right hand. This analysis was conducted at the volume level (voxel-based) but using the outer and inner cortices obtained from Freesurfer as a constraint to select the voxels within each searchlight, as implemented in CoSMoMVPA (Oosterhof et al., 2016; Oosterhof, Wiestler, Downing, & Diedrichsen, 2011). This allowed us to distinguish between regions that are adjacent on the surface (e.g., SI and M1). The results were then projected onto the surface patch enclosing the central voxel. Each searchlight consisted of 100 voxels (the central voxel and its 99 closest neighbors within the ROI). A similar approach has been recently used by Carey, Miquel, Evans, Adank and McGettigan (2017). The searchlight was conducted in the subject space and the resulted maps were then resampled to a common space (fsaverage) for the group analysis.

#### Procedure for the main experiment

The main analysis is described in Figure 4. For each searchlight (100 voxels), we had 9 neural patterns of betas (i.e., one for each of the stimulated locations), from which we computed the 36 pairwise Euclidean distances. We decided to use Euclidean distances rather than correlation coefficients, because it seems a more appropriate measure to adopt in the present context, given that our purpose was to estimate spatial distances on the skin surface. We then used multidimensional scaling to construct a 2-D representational configuration of the skin from this distance matrix, analogous to the way we constructed a perceptual configuration from the matrix of judged distances in the behavioral experiment. The resulting shape was compared with the grid obtained from the behavioral experiment (and to the actual grid on the hand; see Figure S5 in the supplementary material). More specifically, we placed the two configurations into Procrustes alignment and calculated the resulting Procrustes distance (i.e., the dissimilarity in shape of the two configurations). The resulting Procrustes distance and corresponding Procrustes coordinates were assigned to the central voxel of the searchlight. Finally, the brain maps of Procrustes distances and coordinates of each participant were normalized to a common space (fsaverage) and averaged across participants. Note that because the Procrustes distance is a measure of dissimilarity, small numbers indicated that the similarity between the neural and the behavioral shapes was high.

**Figure 4.**
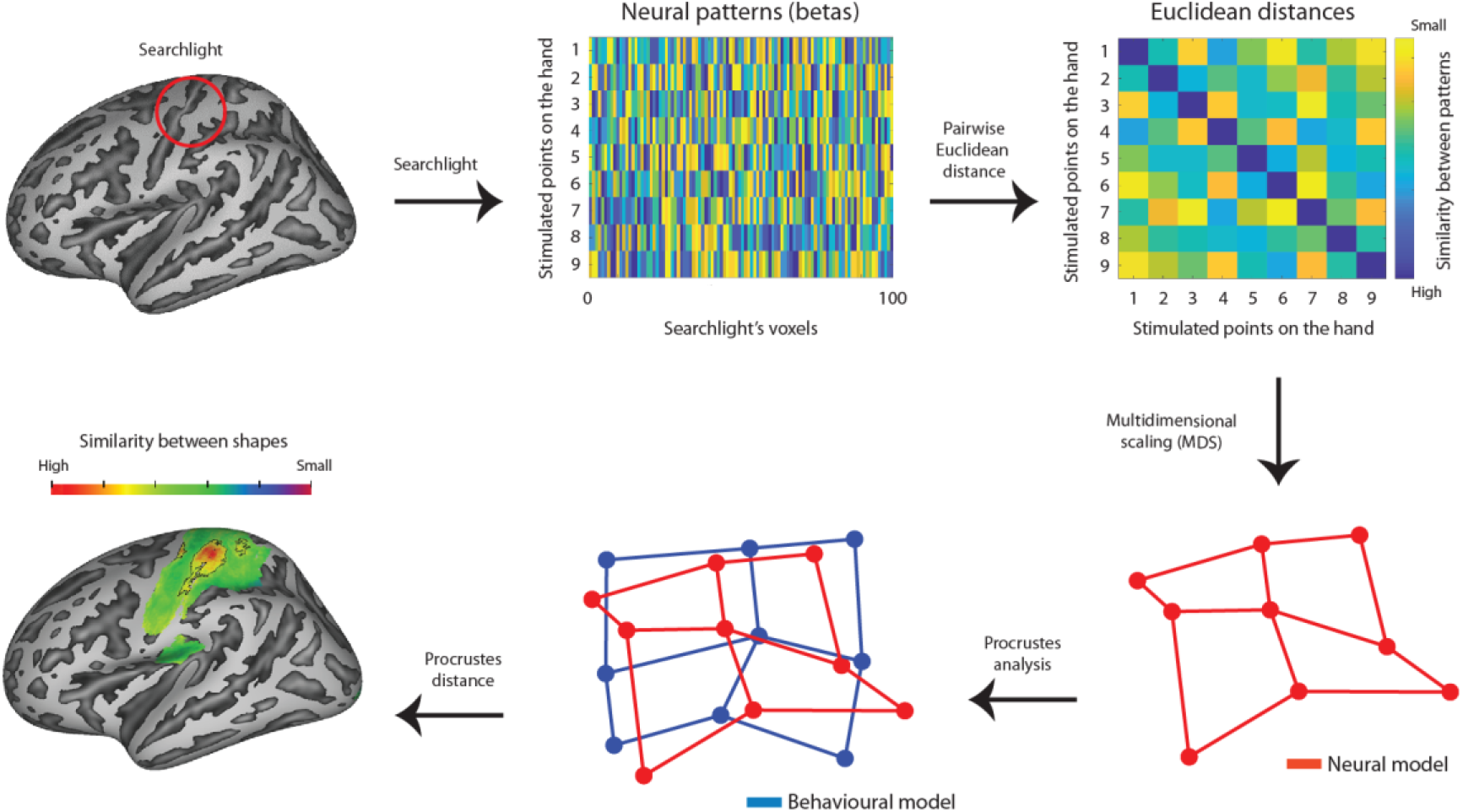
Schematic representation of the analyses steps for the fMRI data which includes: first, we used a searchlight analysis within different ROIs to find activity patterns related to each of the stimulated point on the skin; second, we computed the 36 pairwise Euclidean distances; third, we used the multidimensional scaling to construct a 2-D configuration of the skin from this distance matrix; fourth, the resulting configurations were compared with the perceptual ones using Procrustes alignment resulting in an index of similarity between the configurations (the Procrustes distance). Such a computation was performed for each searchlight and plotted on the brain anatomy. Small Procrustes distances indicate greater similarity between the perceptual and neural configurations.

The rationale for using the behavioral configurations was that these were the only representations that we knew existed in the brain since they were derived from the behavioral data. By contrast, the actual configurations, to the best of our knowledge, could only exist in the physical world. Indeed, it may be that such configurations are not present at all at the neural level. However, in order to assess the potential effect of the actual configurations, the same procedure was performed also using such shapes (see Figure S5 of the supplementary material for a comparison between the actual and behavioral configurations).

To evaluate which of the observed Procrustes distances were statistically smaller than chance, we ran a permutation analysis as described by Stelzer and colleagues (2013) to obtain a threshold size that a cluster (i.e. a set of neighboring vertices) should have in order to be considered statistically significant (with p<0.001 at the vertex level and p<0.05 at the cluster level, as suggested by Stelzer, Chen and Turner, 2013). For each participant, we re-ran the same searchlight analysis as described above, but shuffling the 9 labels before computing the Euclidean distances and we repeated the procedure 100 times. We thus obtained 100 random Procrustes maps for each participant. We then carried out a bootstrap procedure in order to build a null distribution of averaged Procrustes distances: at each iteration, we randomly sampled (with replacement) one map from each participant’s random Procrustes map and we then averaged across these 12 random maps. We repeated this procedure 10,000 times. Then, we computed a p-value at each vertex as the proportion of bootstrap samples that gave a Procrustes distance smaller than the actual Procrustes distance. We thus selected only those vertices that had a p-value smaller than 0.001. Finally, we evaluated the threshold for a cluster to be statistically significant. We individuated neighboring vertices that survived this threshold. We thus obtained a cluster size distribution. To evaluate the p-value associated with each cluster size, we divided the number of clusters for each size by the total number of clusters. The resulting p-values were corrected using a false discovery rate (FDR) of 0.05 and the associated cluster size was used as threshold to select significant clusters in the observed data (see Table 1).

As a final step, we quantified the distortions of neural configurations at representational level within significant clusters adopting the same procedure described for the behavioral study: we extracted the shape associated to each significant cluster by averaging the Procrustes coordinates associated with the vertices within the cluster. Then, we stretched a square grid reflecting the locations of the 9 points by different amounts to find the stretch that minimized the Procrustes distance with each participant’s neural configuration. As for the behavioral data, values between 0.2 and 5 were tested by exhaustive search with a resolution of 0.0005 units in natural logarithm space (i.e., 6,438 steps).

## Acknowledgement

LT, RT and MRL were supported by a grant from the European Research Council (ERC-2013-StG-336050) under the FP7 to MRL.

