## Supplementary_materials for "Reconstructing neural representations of tactile space"

**Supplementary Material**

**Figure S1. Behavioral results**. Generalized Procrustes alignment of the actual configuration of points on the hand (green dots and lines) and perceptual maps (blue dots and lines) for each participant for Experiment 1.

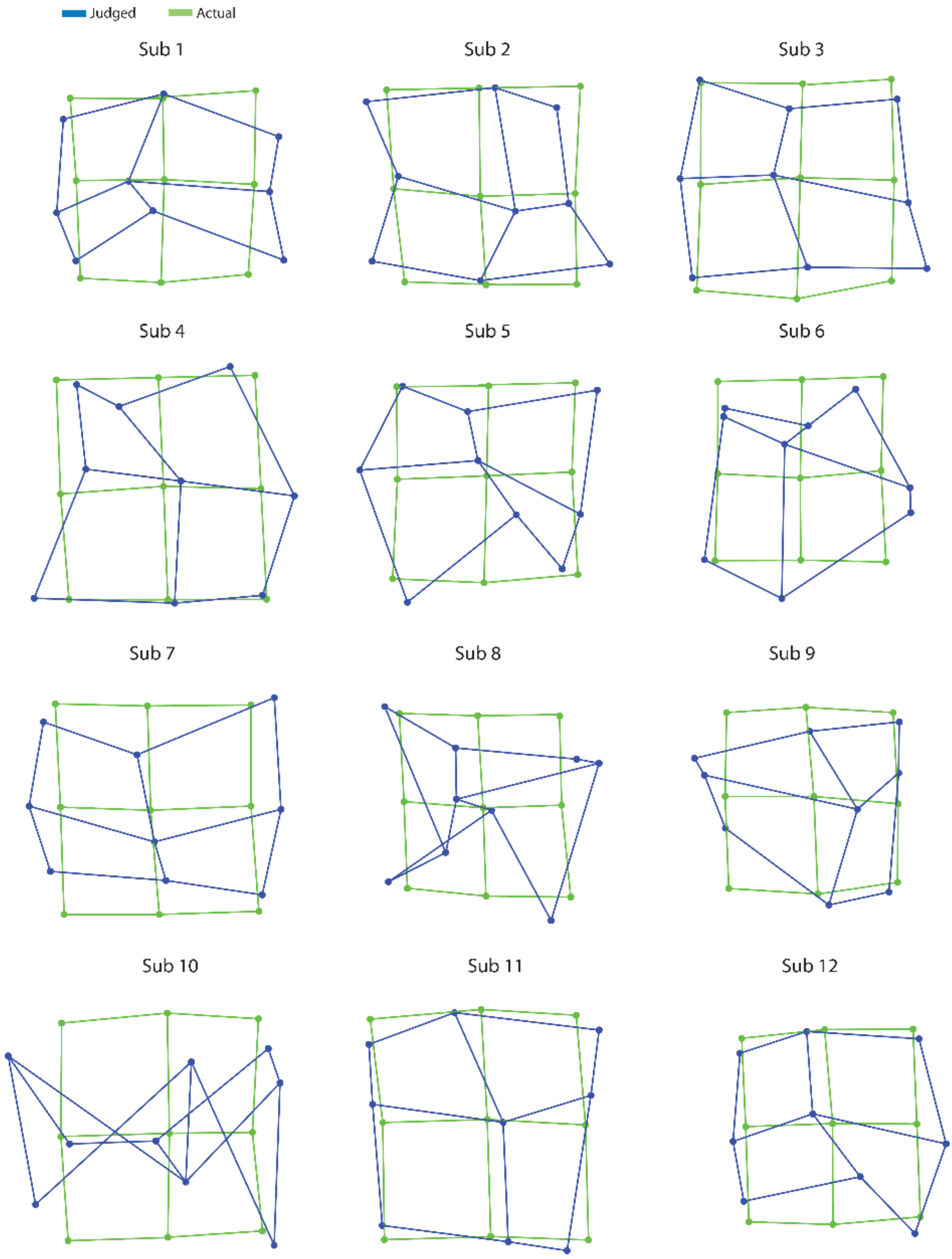

**Figure S2. Individual data.** Mean Procrustes distance between behavioral (blue), fMRI (red) and actual (green) grid on the participants' hand dorsum maps and idealized grids stretched by different amounts in the primary motor cortex (M1: area 4). A stretch of 1 indicates a square grid; stretches greater than 1 indicate stretch in the medio-lateral axis, while stretches less than 1 indicate stretch in the proximo-distal axis. The vertical lines indicate the best-fitting stretches for fmri maps (red) and behavioral maps (blu) for each participant.

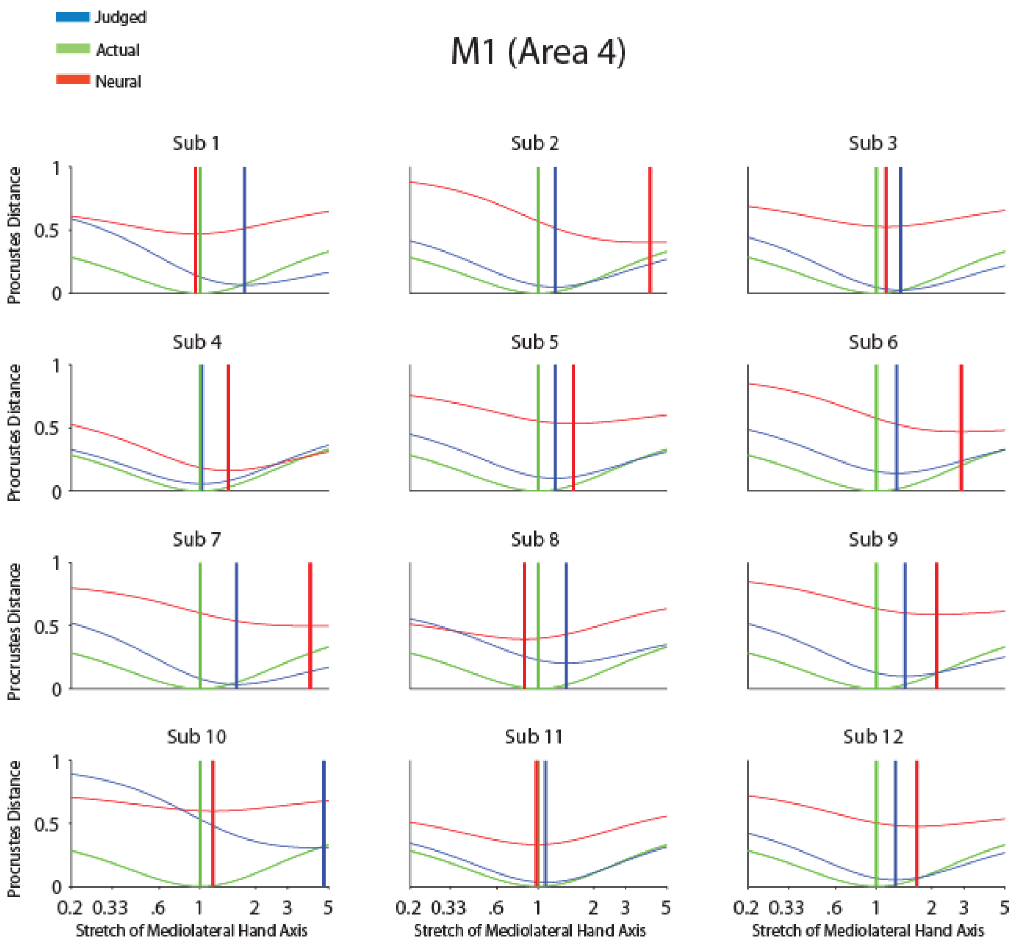

**Figure S3. Individual data.** Mean Procrustes distance between behavioral (blue), fMRI (red) and actual (green) grid on the participants' hand dorsum maps and idealized grids stretched by different amounts in the primary somatosensory cortex (S1: area 3b/1). The vertical lines indicate the best-fitting stretches for fmri maps (red) and behavioral maps (blu) for each participant.

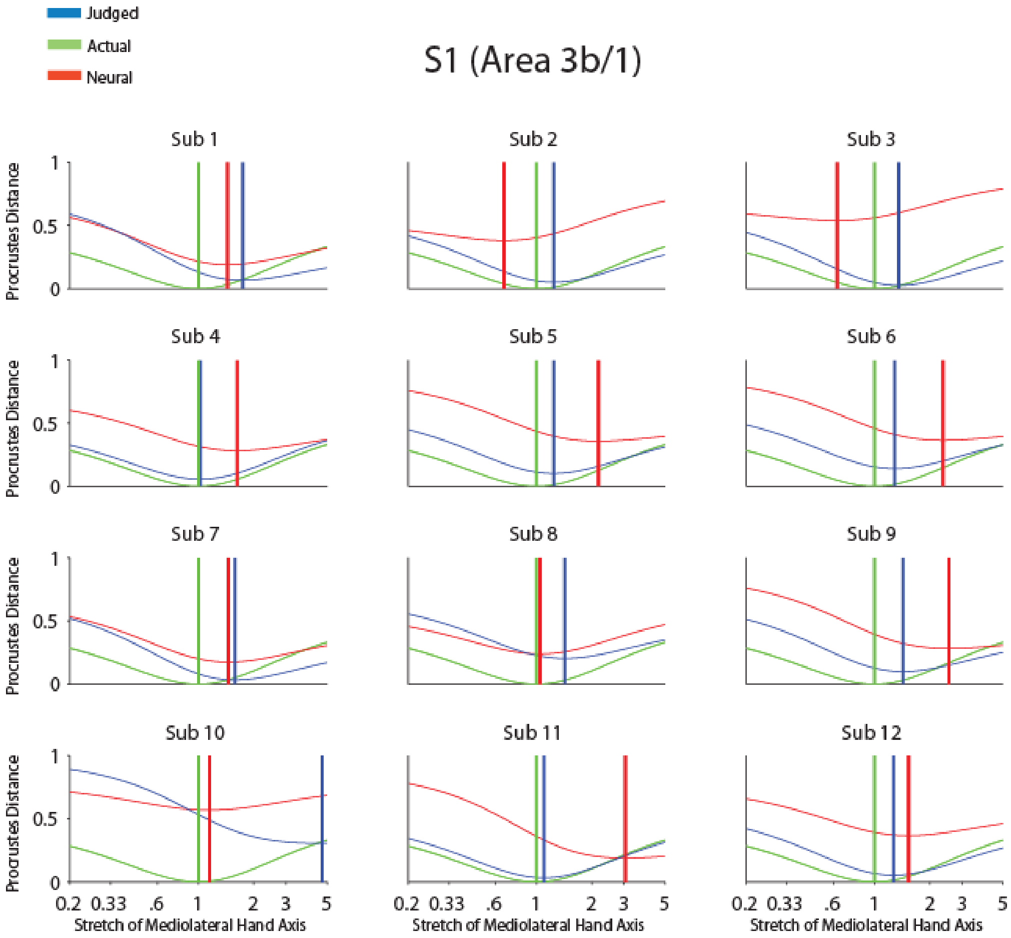

**Figure S4. Individual data.** Mean Procrustes distance between behavioral (blue), fMRI (red) and actual (green) grid on the participants' hand dorsum maps and idealized grids stretched by different amounts in the primary somatosensory cortex (S1: area 2). The vertical lines indicate the best-fitting stretches for fmri maps (red) and behavioral maps (blu) for each participant.

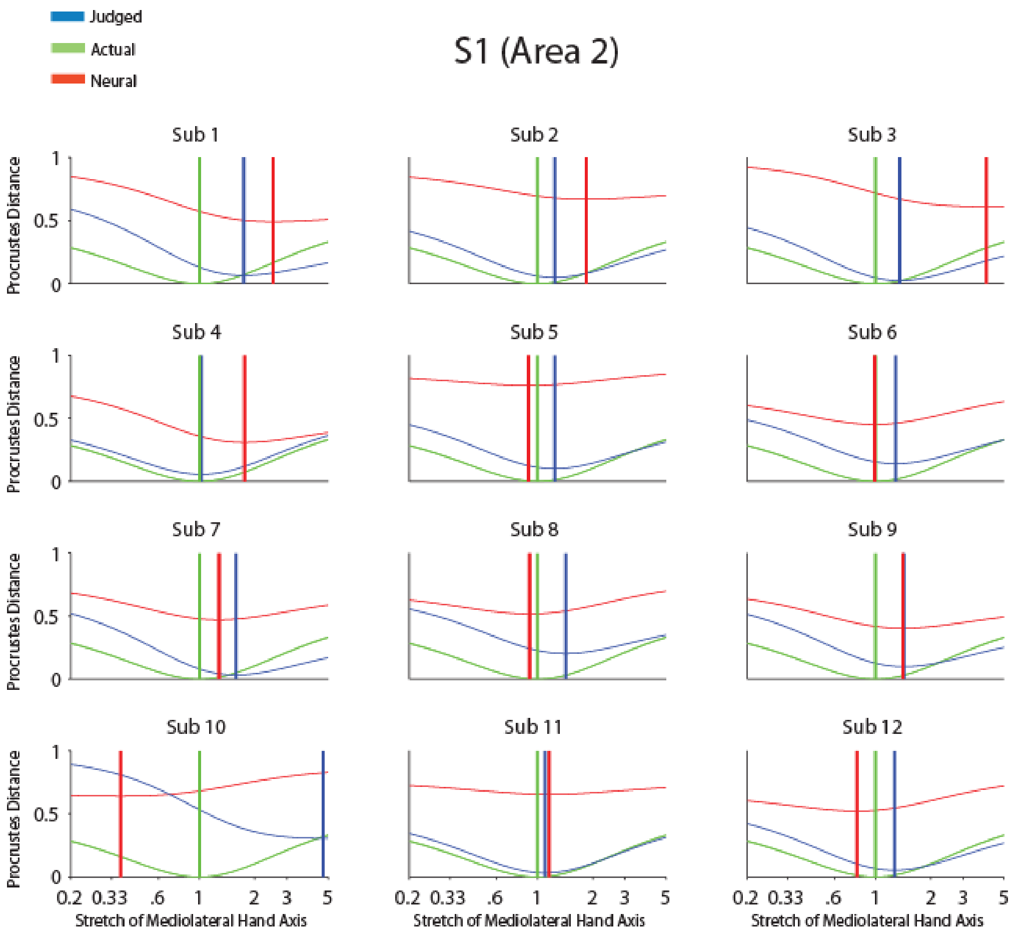

**Figure S5. Averaged data.** Mean Procrustes distance between behavioral (blue), fMRI (red: for each participant) and actual (green) grid on the participants' hand dorsum maps and idealized grids stretched by different amounts when the neural data were placed in Procrustes alignment with the actual (left panel) and behavioral (right panel) maps in different ROIs. As clearly showed by the figures there are no differences in the best-fitting stretches of the activity pattern from the fMRI data when they were placed in Procrustes alignment with the actual or behavioral maps. The shaded regions indicate one standard error of the mean. The vertical lines indicate the mean of the best-fitting stretches for fmri maps (red) and behavioral maps (blu).

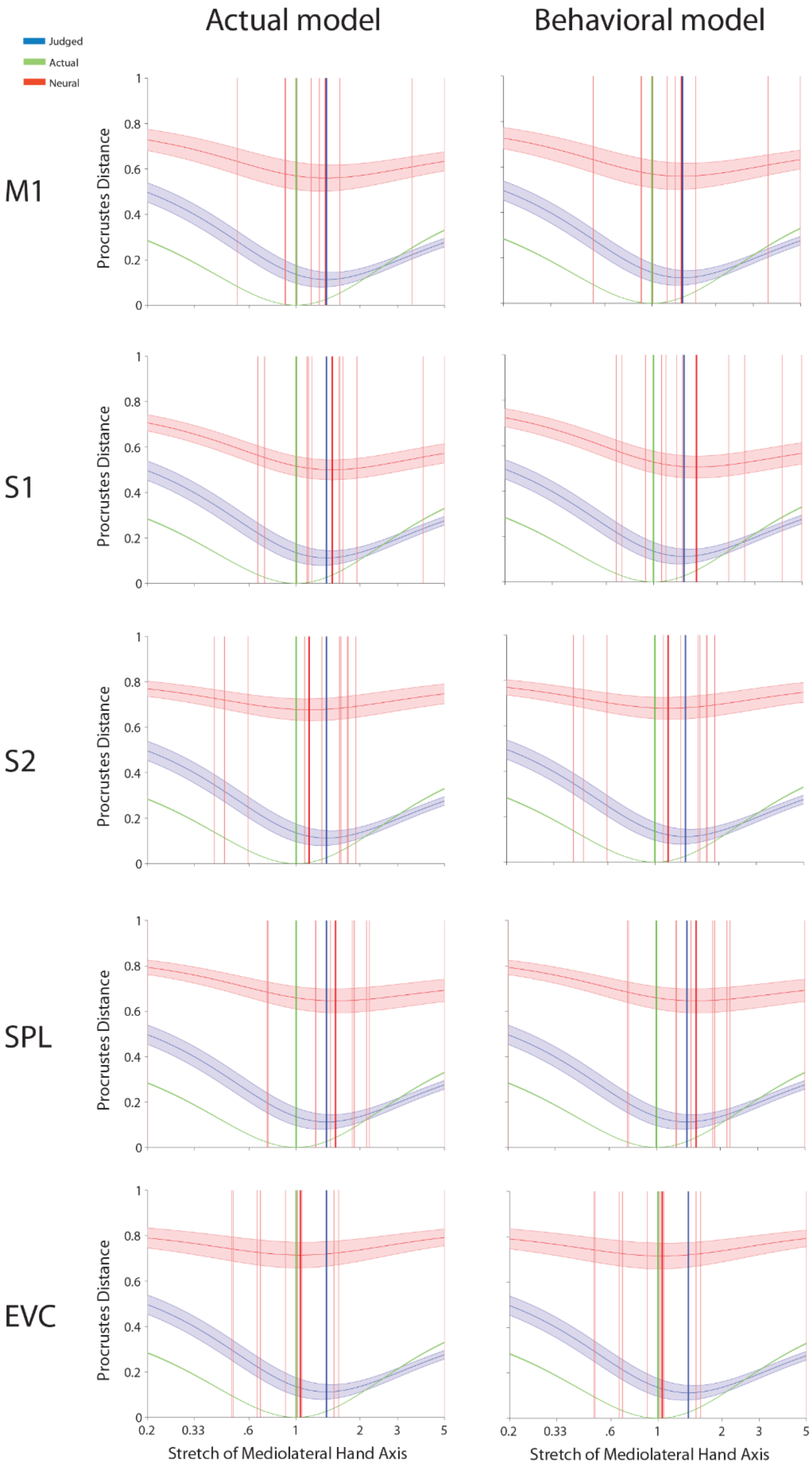

**Figure S6. Statistical analysis.** Probability distribution of cluster size as resulting from cluster-based bootstrapping analysis analysis ($p<0.001$ at the vertex level; FDR<0.05 at the cluster level) in the considered regions of interest. A cluster of the actual data was considered significant if its size (red line) was larger than the cluster size threshold of the ROI (dotted line). This analysis is based on Stelzer, Chen, & Turner (2013) and it is described in the Materials and Method section of the main text.

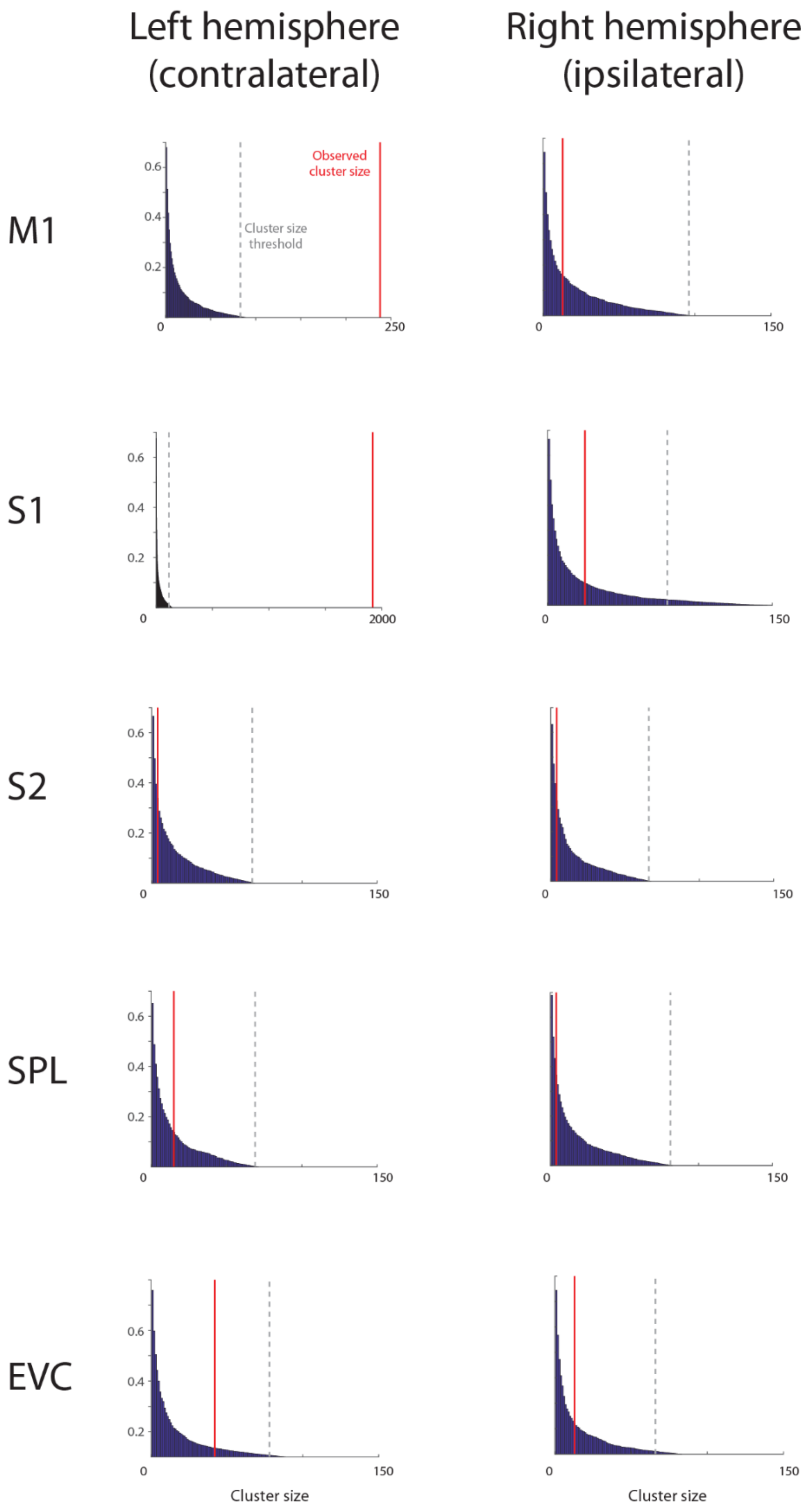

**Figure S7. Statistical analysis.** Significant clusters resulted from the cluster-based bootstrapping analysis ($p<0.001$ at the vertex level; $FDR<0.05$ at the cluster level) on the whole brain. The resulted significant clusters are left Areas 6d/4, 3b/1, 2, OP4, 55b (i.e., contralateral to the locus of stimulation) and right Superior Temporal Visual Area (STV) as well as right Parietal Operculum (i.e., ipsilateral to the locus of stimulation). See **Figure S8** for the related reconstruction of the tactile space.

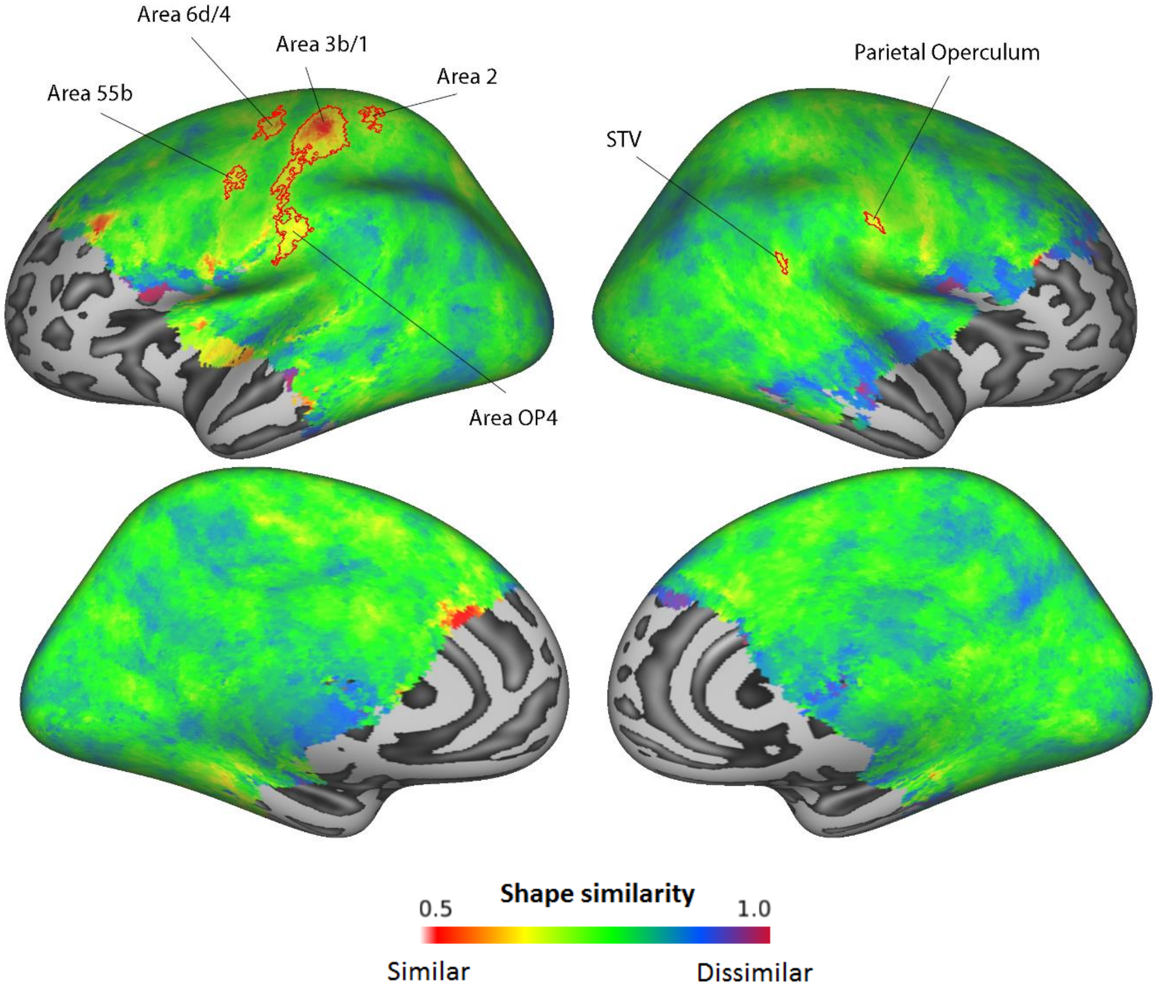

**Figure S8**. **Shape reconstruction.** Reconstruction of the spatial geometry of the skin for the significant clusters resulted from the cluster-based bootstrapping analysis (p<0.001 at the vertex level; FDR<0.05 at the cluster level) at the whole brain level (see **Figure S7**).

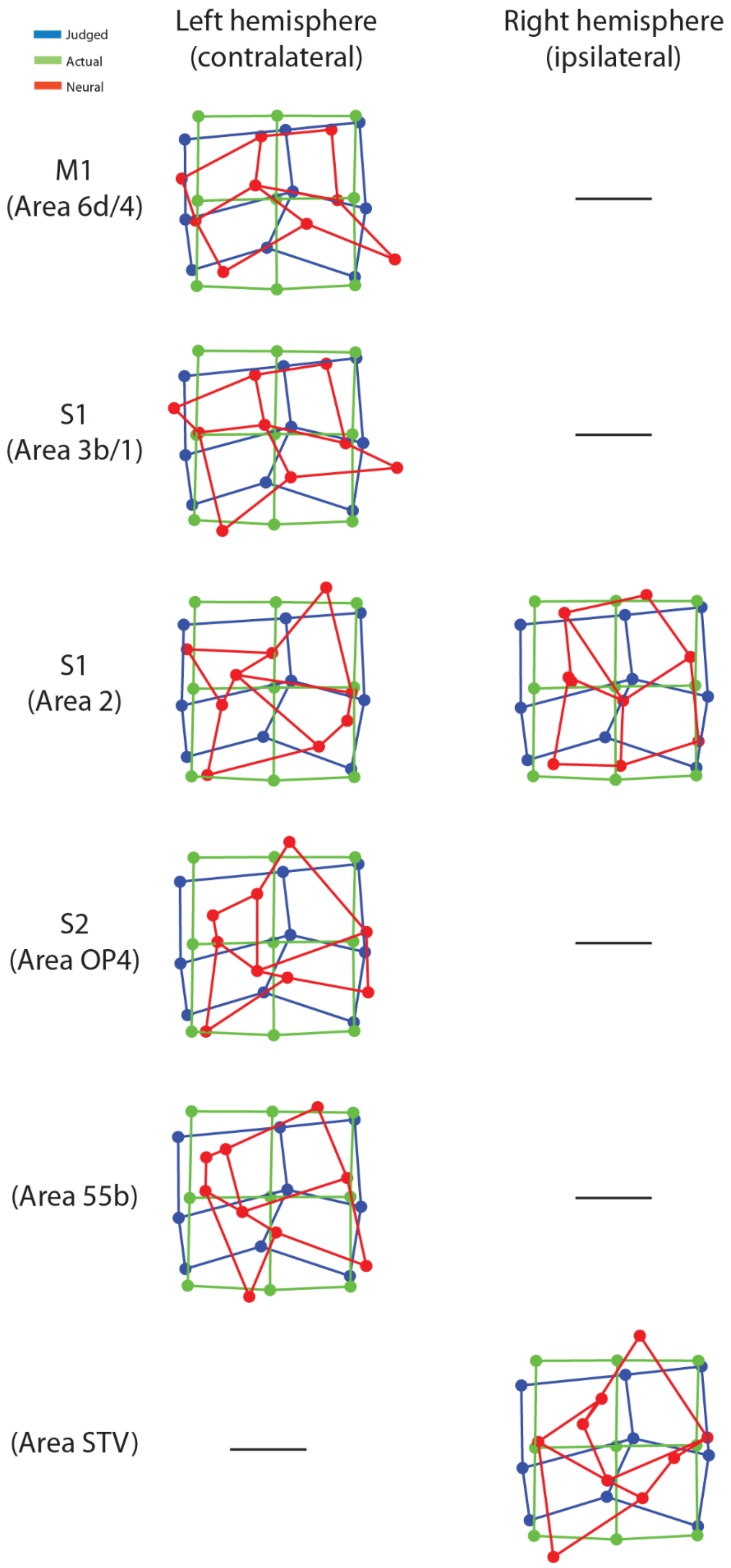
